# The Envelope Proteome Changes Driven by RamA Overproduction in *Klebsiella pneumoniae* that Enhance Acquired β-Lactam Resistance

**DOI:** 10.1101/133918

**Authors:** Juan-Carlos Jiménez-Castellanos, Wan Ahmad Kamil Wan Nur Ismah, Yuiko Takebayashi, Jacqueline Findlay, Thamarai Schneiders, Kate J. Heesom, Matthew B. Avison

## Abstract

**OBJECTIVES:** In *Klebsiella pneumoniae,* overproduction of RamA results in reduced envelope permeability and reduced antimicrobial susceptibility but clinically relevant resistance is rarely observed. Here we have tested whether RamA over-production can enhance acquired β-lactam resistance mechanisms in *K. pneumoniae* and have defined the envelope protein abundance changes seen upon RamA overproduction during growth in low and high osmolarity media.

**METHODS:** Envelope permeability was estimated using a fluorescent dye accumulation assay. Antibiotic susceptibility was measured using disc testing. Total envelope protein production was quantified using LC-MS/MS proteomics and transcript levels quantified by Real Time RT-PCR.

**RESULTS:** RamA overproduction enhanced β-lactamase mediated β-lactam resistance, in some cases dramatically, without altering β-lactamase production. It increased production of efflux pumps and decreased OmpK35 porin production, though *micF* over-expression showed that OmpK35 reduction has little impact on envelope permeability. A survey of *K. pneumoniae* bloodstream isolates revealed *ramA* hyperexpression in 3 out of 4 carbapenemase producers, 1/21 CTX-M producers and 2/19 strains not carrying CTX-M or carbapenemases.

**CONCLUSIONS:** Whilst RamA is not a key mediator of antibiotic resistance in *K. pneumoniae* on its own, it is potentially important for enhancing the spectrum of acquired β-lactamase mediated β-lactam resistance. LC-MS/MS proteomics analysis has revealed that this enhancement is achieved predominantly through activation of efflux pump production.

## Introduction

RamA is a global transcriptional activator ^1^ found in, amongst other Enterobacteriaceae: *Salmonella* spp.,^2^ *Enterobacter* spp.^1,3^ and *Klebsiella* spp.^1,4^ but not *Escherichia coli.* Where characterised, RamA has revealed a function comparable to *E. coli* MarA.^1-5^ In wild-type *Klebsiella pneumoniae,* at least under standard laboratory growth conditions, production of RamA is low because of RamR, a transcriptional repressor that occludes *the ramA* promoter.^6^ In some clinical isolates, RamA is overproduced due to de-repressing mutations in *ramR*.^7-10^ RamA activates the transcription of a regulon including *oqxAB* and *acrAB,* encoding the components of two tripartite antimicrobial drug efflux pumps, and *tolC,* which encodes the outer membrane protein used by both.^4^

Overexpression of *ramA* in *K. pneumoniae* isolates that lack other antibiotic resistance mechanisms increases MICs of a wide range of antimicrobials, including cephalosporins but not carbapenems. However, even overexpressing *ramA* >1000 fold only confers clinically relevant resistance to one or two antimicrobials and only in some isolates.^5^ On its own, therefore, RamA is not a key resistance determinant in *K. pneumoniae,* however RamA over-producing clinical isolates can carry acquired resistance mechanisms, particularly plasmid encoded β-lactamases.^11^ Accordingly, the first aim of the work presented here was to determine whether RamA overproduction can enhance the spectrum of resistance conferred by acquired β-lactamases. We also wanted to test whether *ramR* loss of function mutations are more common in cephalosporinase or carbapenemase producing *K. pneumoniae* clinical isolates than in isolates that do not carry these types of enzymes.

We were particularly keen to investigate whether RamA-mediated reduced carbapenem susceptibility occurs in isolates producing ESBLs or AmpC type cephalosporinases. The rationale underlying this aim was based on our previous observation of RamA-mediated OmpK35 downregulation,^4,5^ as loss of function mutations in OmpK35 have previously been shown to increase carbapenem MICs against *K. pneumoniae* isolates carrying ESBLs or AmpC β-lactamases.^12^

The enhancement of β-lactam MICs seen following OmpK35 loss of function in a cephalosporinase producing *K. pneumoniae* is reportedly minimised during growth in high osmolarity Muller Hinton media, which is the medium of choice for most antibiotic susceptibility testing protocols,^13^ but is maximised during growth in low osmolarity Nutrient media. This is because these media reportedly support different basal OmpK35 levels, as defined using outer membrane protein profiling and SDS-PAGE.^12^ Accordingly, we also set out to define the envelope proteome changes stimulated by RamA overproduction in Muller Hinton broth and Nutrient broth using a much more discriminatory and accurate methodology: Orbitrap liquid chromatography tandem mass spectrometry (LC-MS/MS). The aim was to identify common and growth medium-specific effects of RamA overproduction, and to confirm a previous report that basal OmpK35 levels are different in the two media.^12^ Finally, we set out to define the contribution of RamA-mediated OmpK35 downregulation to the overall effect of RamA overproduction on envelope permeability and antibiotic susceptibility in *K. pneumoniae.*

## Materials and Methods

### Bacterial strains and antibiotic susceptibility testing

*E. coli* TOP10 (Invitrogen, Leek, The Netherlands), 44 non-replicate *K. pneumoniae* human bloodstream isolates having various antimicrobial susceptibility profiles (provided by Dr Karen Bowker, Department of Microbiology, North Bristol NHS Trust), *K. pneumoniae* NCTC5055 transformants carrying pBAD(*ramA*) or a pBAD vector only control plasmid ^5^ and the otherwise isogenic pair ECL8 ^1^ and ECL8Δ*ramR* ^4^ were used throughout. Disc susceptibility testing was performed according to CLSI methodology ^13^ and interpreted using CLSI performance standards.^14^

### Cloning plasmid-mediated β-lactamase genes and micF and sequencing ramR

Cloning *bla*_NDM-1_ (with its IS*Abal25* promoter) into the cloning vector pSU18 ^15^ has previously been reported.^16^ *K. pneumoniae micF* and the following additional β-lactamase genes were synthesised or amplified by PCR from the sources and using the primers listed in Table SI and the PCR method previously described ^17^ in such a way as to include their native promoters: *bla*_IMP-1_ and *bla*_VIM-1_ (with hybrid and weak strength class 1 integron promoters, respectively),^18^ *bla*_CTX-M1_ and *bla*_CMY-4_ (with IS*Ecp1* promoters), *bla*_KPC-3_ (with IS*Kpn7* promoter) and *bla*_OXA-48_ (with IS*1999* promoter). Some PCR amplicons were TA cloned into the pCR2.1-TOPO cloning vector (Invitrogen) according to the manufacturer’s instructions. These pCR2.1 inserts, removed by restriction enzyme digestion, and other PCR amplicons directly cut with restriction enzymes, were ligated into the pSU18 ^15^ or pK18 ^19^ cloning vectors as illustrated in Table SI. Plasmids were used to transform *K. pneumoniae* isolates to chloramphenicol or kanamycin (30 mg/L) resistance for pSU18 and pK18, respectively, using electroporation as standard for lab strain *E.coli.* To sequence *ramR* from clinical isolates, the gene was first amplified by PCR as previously ^17^ and sequenced using the primers recorded in Table S1. Sequence alignment and analysis were performed using the online ClustalOmega (http://www.ebi.ac.uk/Tools/msa/clustalo/) multiple sequence alignment program to determine the *ramR* mutations in the isolates relative to the reference sequence *K. pneumoniae* Ecl8 (Accession Number HF536482.1).

### Growth of cultures for all experiments

Each strain or transformant was inoculated into a separate batch of 50 mL Cation Adjusted Muller-Hinton Broth (Sigma) or Nutrient Broth (Oxoid) containing appropriate selection in a 250 mL foam stoppered flask to an initial Optical Density at 600 nm (OD_600_) of ≈0.05. Cultures were incubated with shaking (160 rpm) until the OD_600_ had reached 0.5-0.7.

### Fluorescent Hoechst (H) 33342 dye accumulation assay

Envelope permeability was estimated as described previously ^5^ in bacteria grown in liquid culture using an established fluorescent dye accumulation assay^20^ with black flat-bottomed 96-well plates (Greiner Bio-one, Stonehouse, UK) and a Fluostar Optima (Aylesbury, UK) plate reader. H33342 (Sigma) was used at a final concentration of 2.5 μM.

### Characterisation of Envelope Proteomes and Real-Time RT-PCR

Methods used for protein and RNA extraction and analysis of abundance were almost identical to those used previously ^5^ and are presented in detail in Supplementary Material. For each LC-MS/MS proteomics experiment, raw protein abundance data were collected for three biological replicates of each growth condition. The significance of any observed difference between the means of the triplicate abundance data for one protein in two different growth conditions was calculated using a T-test comparing the raw abundance data as described in Supplementary Material.

### β-Lactamase assays and multiplex PCR for β-lactamase gene carriage

Multiplex PCRs were performed for (i) carbapenemase genes encoding the IMP, VIM, OXA-48-like, NDM and KPC enzymes,^21^ though the NDM and OXA-48 like gene identifying PCRs were run separately and not as a multiplex with the rest (ii) genes encoding CTX-M groups 1, 2, 8, 9 and 25 ^22^ (iii) a bespoke multiplex for genes encoding TEM-1, SHV, OXA-1 and CMY. All multiplex PCR primers and expected product sizes are recorded in Table S2. DNA template was prepared as previously described ^17^ and 1 μL of supernatant used in a final PCR reaction volume of 25 μL consisting of 12.5 μL REDTaq ReadyMix (Sigma) with 10 μM of each primer. PCR was performed using the following conditions: 94°C for 10 min, 35 cycles of amplification consisting of 94°C for 1 min, 52-56°C for 1 min, 72°C for 1 min 30 s and a final extension at 72°C for 10 min.

β-Lactamase assays were performed as follows: overnight cultures of bacteria were diluted to an optical density at 600 nm (OD_600_) of 0.1 in Muller-Hinton broth and grown at 37°C until the OD_600_ was 0.8 before cell extracts were prepared and levels of β-lactamase activity in cell extracts measured as described previously^23^ using 100 μM nitrocefin as a substrate. Linear gradients (ΔAU/min) were extrapolated and an extinction coefficient of 17,400 AU/M was used to calculate nitrocefin hydrolysing activity. The total protein concentration in each cell extract was quantified using the Bio-Rad protein assay reagent (Bio-Rad, Hemel Hempstead, United Kingdom) according to the manufacturer’s instructions and used to calculate relative specific enzyme activity (pmol/min/μg) in each cell extract.

## Results and Discussion

### Enhancement of β-lactam resistance in K. pneumoniae by RamA in the presence of acquired β-lactamases

A library of seven clinically important mobile β-lactamase genes was created using pSU18, a broad host-range, low copy number vector.^15^ Each gene was expressed from a native promoter commonly seen in clinical isolates to give close to wild-type levels of β-lactamase production (see Table SI). Once used to transform *K. pneumoniae* Ecl8 and Ecl8Δ*ramR*,^4,5^ this library allowed us to compare the relative importance of these mobile β-lactamase genes and the additional impact of overproducing RamA on β-lactam resistance. It is important to note that there was no significant change in β-lactamase production in Ecl8Δ*ramR* transformants versus Ecl8 transformants carrying the same β-lactamase gene (Table S3)

Ecl8 is susceptible to 17/18 tested β-lactams (Table 1). Overproduction of RamA (i.e. Ecl8Δ*ramR)* reduced zone diameters ≥2 mm for 13/18 tested β-lactams, but clinically relevant non-susceptibility (resistance or intermediate resistance based on CLSI breakpoints)^14^ was not achieved for any of them (Table 1). Introduction of β-lactamase genes into Ecl8 had significant effects on β-lactam susceptibility, as expected. The metallo-β-lactamases (MBLs) NDM-1 and IMβ-1 rendered Ecl8 non-susceptible to 17/18 and 14/18 β-lactams, respectively. Because of this, it was difficult to see any additional effect that might be conferred by RamA overproduction (Table 1). Accordingly, the gene encoding VIM-1 was deliberately cloned with a weak integron promoter and the resultant low-level MBL production meant that Ecl8 remained susceptible to 12/18 test β-lactams. In this case, the effect of RamA overproduction was profound. Ecl8Δ*ramR* (VIM-1) was only susceptible to aztreonam and cefotetan; importantly, 10 β-lactams were rendered ineffective by RamA overproduction in the presence of low-level MBL activity (Table 1).

**Table 1.**

|  | pSU18 |  | IMP-1 |  | NDM-1 |  | VIM-1 |  | CTX-M1 |  | CMY-4 |  | KPC-3 |  | OXA-48 |  |
| --- | --- | --- | --- | --- | --- | --- | --- | --- | --- | --- | --- | --- | --- | --- | --- | --- |
| Antibiotic (μg) | Ecl8 | Δ | Ecl8 | Δ | Ecl8 | Δ | Ecl8 | Δ | Ecl8 | Δ | Ecl8 | Δ | Ecl8 | Δ | Ecl8 | Δ |
| Ampicillin (10) | 16 | 10 | 6 | 6 | 6 | 6 | 6 | 6 | 11 | 6 | 6 | 6 | 6 | 6 | 12 | 6 |
| Piperacillin (100) | 29 | 25 | 27 | 22 | 9 | 6 | 22 | 15 | 15 | 6 | 17 | 14 | 8 | 6 | 28 | 26 |
| Piperacillin/<br>Tazobactam (110) | 32 | 26 | 30 | 22 | 8 | 6 | 22 | 15 | 30 | 25 | 25 | 20 | 16 | 8 | 30 | 28 |
| Cefotetan (30) | 32 | 30 | 15 | 6 | 14 | 6 | 30 | 22 | 30 | 28 | 17 | 14 | 21 | 13 | 31 | 27 |
| Cefoxitin (30) | 30 | 23 | 6 | 6 | 6 | 6 | 22 | 15 | 28 | 23 | 11 | 6 | 23 | 10 | 28 | 20 |
| Cefuroxime (30) | 32 | 22 | 6 | 6 | 6 | 6 | 12 | 6 | 16 | 6 | 6 | 6 | 6 | 6 | 30 | 21 |
| Cefotaxime (30) | 40 | 32 | 20 | 12 | 6 | 6 | 22 | 15 | 15 | 6 | 15 | 12 | 19 | 10 | 39 | 32 |
| Ceftriaxone (30) | 38 | 33 | 19 | 14 | 6 | 6 | 25 | 16 | 16 | 6 | 15 | 10 | 13 | 6 | 35 | 32 |
| Ceftazidime (30) | 33 | 30 | 15 | 6 | 6 | 6 | 19 | 11 | 20 | 15 | 13 | 7 | 14 | 6 | 31 | 29 |
| Cefoperazone (75) | 34 | 29 | 19 | 15 | 10 | 6 | 22 | 15 | 17 | 6 | 22 | 20 | 11 | 8 | 30 | 26 |
| Ceftizoxime (30) | 40 | 35 | 20 | 15 | 6 | 6 | 25 | 19 | 30 | 15 | 18 | 16 | 20 | 11 | 40 | 33 |
| Cefixime (5) | 33 | 29 | 14 | 6 | 6 | 6 | 18 | 6 | 16 | 6 | 6 | 6 | 17 | 6 | 31 | 27 |
| Cefepime (30) | 36 | 36 | 25 | 20 | 18 | 10 | 26 | 20 | 22 | 15 | 37 | 33 | 19 | 11 | 35 | 31 |
| Aztreonam (30) | 40 | 38 | 40 | 37 | 40 | 38 | 40 | 37 | 19 | 15 | 18 | 15 | 12 | 6 | 38 | 33 |
| Doripenem (10) | 30 | 30 | 20 | 18 | 15 | 10 | 23 | 21 | 30 | 30 | 26 | 26 | 16 | 16 | 28 | 28 |
| Ertapenem (10) | 30 | 30 | 20 | 15 | 10 | 6 | 26 | 20 | 29 | 22 | 23 | 20 | 17 | 9 | 24 | 22 |
| Imipenem (10) | 30 | 30 | 16 | 16 | 14 | 10 | 19 | 19 | 30 | 30 | 30 | 30 | 14 | 11 | 26 | 25 |
| Meropenem (10) | 30 | 30 | 18 | 15 | 8 | 8 | 23 | 20 | 30 | 30 | 31 | 31 | 18 | 12 | 30 | 27 |

Assays were performed using Muller Hinton agar according to the CLSI protocol<sup>13</sup> for *K. pneumoniae* Ecl8 and Ecl8ΔramR (Δ). Values reported are the means of three repetitions rounded to the nearest integer. Zones confirming non-susceptibility are shaded grey. Susceptibility breakpoints are as set by the CLSI.<sup>14</sup>

The possibility that reduced envelope permeability can enhance the ability of serine active site β-lactamases (SBLs) such as CTX-M or AmpC type cephalosporinases to confer carbapenem resistance in *K. pneumoniae* has previously been suggested.^12^ We found evidence that this is correct, but only for ertapenem. Ecl8 remained susceptible to β-lactam/β-lactamase inhibitor combinations, cephamycins and carbapenems when carrying CTX-M1. Susceptibility to all these β-lactam classes was retained following overproduction of RamA. There was a considerable zone diameter reduction for ertapenem (by 7 mm; on the verge of being non-susceptible) but for the other carbapenems, no change at all. In the case of the AmpC-type SBL CMY-4, ertapenem resistance was conferred when RamA was overproduced, though the other carbapenems remained equally active against Ecl8Δ*ramR* (CMY-4) as against Ecl8 (CMY-4). Finally, we tested SBLs with carbapenemase activity. In Ecl8 carrying KPC-3, only the cephamycins retained activity, but even these were lost following overproduction of RamA, underlining the threat of this broad-spectrum enzyme. In contrast, the effect of carrying OXA-48 on β-lactam resistance was relatively weak. When RamA was overproduced there was some inhibition zone diameter reduction, particularly for the carbapenems, but Ecl8Δ*ramR*(OXA-48) remained susceptible to 17/18 β-lactams (Table 1).

### A survey of ramR function in clinical isolates carrying or not carrying cephalosporinases and carbapenemases

In order to perform a survey of *ramR* mutation in *K. pneumoniae,* 44 bloodstream isolates were randomly collected. PCR was used to determine *ramR* sequence in these isolates. Nine isolates had mutations in *ramR,* (in comparison with Ecl8), in addition to a mutation encoding the Thr141Ile variant, seen in several isolates, which is known to arise via genetic drift and has no impact on phenotype.^9^ Two of the isolates carried the same *ramR* mutation, encoding a Ala19Val variant. To confirm which of the eight different mutations reduce the repressor function of RamR, and therefore caused hyperexpression of *ramA,* we used quantitative Real–Time RT-PCR to measure *ramA* transcript levels in comparison with the *ramR* wild-type isolate AE. Six of the mutants hyper-expressed *ramA* with a range of 4.3-fold to 70.6-fold more than in wild-type clinical isolate AE (Table S4).

To see if there was any linkage between *ramR* mutation and carriage of β-lactamase genes, three multiplex PCRs were performed to categorise the β-lactamase genes present in each of the 44 bloodstream isolates. Four carried at least one carbapenemase gene, of which two also carried *bla*_CTX-M_; 21 additional isolates carried *bla*_CTX-M_. All CTX-M genes were of group 1. The remaining 19 isolates did not carry any cephalosporinase or carbapenemase gene (Table S4).

Finally, disc susceptibility testing was performed for eight β-lactams against the 44 clinical isolates. Using the sum of all the inhibition zone diameters to represent the combined β-lactam susceptibility for each isolate, we ranked the isolates (Table S4). Not surprisingly the four carbapenemase gene positive isolates were the four least β-lactam susceptible isolates overall. Nonetheless, it was striking to find that 3 out of 4 of these isolates also hyperexpress *ramA* because of a *ramR* mutation. Only one of the 21 carbapenemase negative, CTX-M positive isolates hyper-expressed *ramA;* but this isolate (isolate T) was by far the least susceptible isolate in this group. As predicted from our transformation experiments, loss of RamR repressor activity in a CTX-M positive background (isolate T) reduced carbapenem susceptibility relative to the 20 isolates that have CTX-M but an intact *ramR.* However, this effect (Table S4) was greater than that seen in the Ecl8/Ecl8Δ*ramR* transformants (Table 1), with resistance to ertapenem, and intermediate resistance to doripenem being observed in isolate T, together with reduced susceptibility to meropenem. There must be some additional mechanism at play in isolate T not seen in Ecl8. As seen with Ecl8Δ*ramR* (Table 1) loss of RamR repressor activity in the two clinical isolates that lack any cephalosporinase or carbapenemase genes had minimal impact on β-lactam susceptibility in comparison with the *ramR* wild-type isolates (Table S4)

### Envelope Proteome Changes Following RamA Overproduction in K. pneumoniae in NB and MHB and Impact on Envelope Permeability

We have recently shown using LC-MS/MS proteomics that RamA overproduction in *K. pneumoniae* NCTC5055 using a pBAD expression plasmid increased the production of two efflux pumps: AcrAB-TolC and OqxAB-TolC.^5^ To obtain a more detailed understanding of the RamA regulon, we used Orbitrap LC-MS/MS proteomics to generate a comprehensive list of envelope proteins whose production is altered upon RamA overproduction. Table S5 shows summary data for the *K. pneumoniae* Ecl8/Ecl8Δ*ramR* otherwise isogenic pair during growth in Muller-Hinton Broth (MHB). It has previously been reported that low osmolarity media such as nutrient broth (NB) affects the levels of OmpK35 relative to growth in high osmolarity media such as MHB.^12^ This observation is potentially important because OmpK35 levels might impact on envelope permeability and because antimicrobial susceptibility assays are generally performed using Muller-Hinton media. Accordingly, to see whether medium osmolarity affects the impact of RamA overproduction and to quantify the effect of medium choice on OmpK35 levels, we also compared envelope proteome changes in the Ecl8/Ecl8Δ*ramR* pair during growth in NB (Table S6).

Our LC-MS/MS methodology allowed identification and absolute quantification with a high degree of certainty (≥ 3 peptides identified) of 655 and 494 proteins in envelope preparations from cells grown in MHB and NB, respectively (see supplementary proteomics data file). Previous data from SDS-PAGE analysis of outer membrane protein preparations have been interpreted as meaning that OmpK35 levels increase upon shifting from MHB to NB and that OmpK36 levels remain constant.^12^ However, because the same amount of total protein was loaded in each SDS-PAGE well, it is not possible to quantify either OmpK35 or OmpK36 in absolute terms, only the ratio of the two can be estimated. Our LC-MS/MS data did confirm this earlier finding ^12^ that there is a shift in the OmpK36:OmpK35 ratio in Ecl8 from 1.7:1 (calculated from mean protein abundance data, *p*=0.013, *n*=3) during growth in MHB to 1.1:1 (*p*=0.45, *n*=3) during growth in NB. However, our absolute abundance data revealed that this change in OmpK36:OmpK35 ratio is achieved through downregulation of OmpK36 (2.2-fold, *p*=0.011, *n*=3) not upregulation of OmpK35 (0.7-fold, *p*=0.16, *n*=3) during growth in NB relative to growth in MHB (see supplementary proteomics data file).

Twenty-nine proteins were found to be ≥ 2-fold up or down regulated in Ecl8Δ*ramR* versus Ecl8 following application of our statistical significance cut-off (p<0.05 for a T-test comparing absolute protein abundance data, *n*=3) during growth in MHB, and 33 proteins were differentially regulated during growth in NB (Tables S5, S6 and additional supplementary proteomics data file). Of these, 12 proteins were similarly regulated in both media; 11 upregulated, with only the OmpK35 porin being downregulated (Table 2). Ten out of 11 proteins upregulated in Ecl8Δ*ramR* in both media were also upregulated in *K. pneumoniae* strain NCTC5055 following overproduction of RamA via the pBAD expression plasmid (Table S7). These 10 core upregulated proteins represent two efflux pumps (AcrAB and OqxAB, together with the outer membrane efflux protein TolC). The remaining proteins are poorly characterised and their precise role in RamA mediated permeability change is currently under investigation.

**Table 2.** Significant Changes in Envelope Protein Abundance Seen in Ecl8Δ*ramR* versus Ecl8 During Growth in Both MHB and NB

| Accession | Description | MHB |  | NB |  |
| --- | --- | --- | --- | --- | --- |
|  |  | *p* value | Fold Change | *p* value | Fold Change |
| A6T5M4 | AcrB | 0.038 | 4.87 | 0.044 | 6.21 |
| A6T5M5 | AcrA | 0.002 | 3.88 | 0.002 | 3.38 |
| A6T5Q5 | Putative Outer Membrane Protein | <0.001 | 5.85 | <0.001 | 20.00 |
| A6T6W3 | Putative lipoprotein YbjP | 0.001 | 5.30 | 0.002 | 7.01 |
| A6T721 | OmpK35 | 0.007 | 0.43 | 0.026 | 0.23 |
| A6T7Z9 | ABC Transporter SapF | <0.001 | >20 | <0.001 | 20.00 |
| A6T8F9 | Heat shock protein HslJ | <0.001 | >20 | <0.001 | 20.00 |
| A6T9Y7 | Putative uncharacterized protein YdhA | 0.021 | 6.17 | <0.001 | 20.00 |
| A6TCQ4 | OqxA | <0.001 | >20 | <0.001 | 20.00 |
| A6TCQ5 | OqxB | <0.001 | >20 | <0.001 | 20.00 |
| A6TCT2 | Putative Uncharacterized Protein | 0.012 | 4.06 | <0.001 | 5.03 |
| A6TE24 | TolC | 0.004 | 3.15 | 0.013 | 3.51 |

Methodology and details of data analysis are described in Supplementary Material

Of the medium-specific impacts of RamA overproduction, two are striking. In MHB only, downregulation of several proteins encoded by the maltose transport operon occurs (Table S5). The LamB2 porin is amongst these. Interestingly, its loss by mutation has been implicated in reduced carbapenem entry in *K. pneumoniae,*^25,26^ so it is conceivable that downregulation of LamB2 might enhance the impact of RamA overproduction on carbapenem MICs during growth on Muller-Hinton agar, which is apparent in the presence of certain plasmid-mediated β-lactamases (Table 1, Table S4). In NB only, the efflux pump AcrEF is upregulated in Ecl8Δ*ramR* relative to Ecl8. Whilst this pump has not been specifically characterised in *K. pneumoniae,* its equivalent has a role in antimicrobial resistance in other enteric bacteria, and it is part of the *Salmonella* RamA regulon.^2^ Also upregulated in NB only is the transporter complex proteins YrbCDEF (Table S6). YrbB, which is encoded in the same operon was below the level of detection by the LC-MS/MS instrument, so we cannot say whether it was upregulated or not, but it seems a likely scenario. RamA-mediated regulation of the *yrb* locus has previously been demonstrated using transcriptome analyses in both *K. pneumoniae* and *S*. *typhimurium* and a potential RamA binding site has been proposed for the *yrb* locus in *K. pneumoniae*.^4^ The four Yrb proteins upregulated in Ecl8Δ*ramR* have >90% identity to the MlaCDEF proteins, part of the MlaABCDEF ABC transporter from *E. coli,* which has recently been shown to play an important role in retrograde phospholipid trafficking. The perturbation of the phospholipid content of the outer membrane is likely to result in a reduction in the sensitivity of the outer membrane to chemical damage, and may also affect antimicrobial/cell affinity, reducing rate of entry.^27-29^ The AcrEF and Yrb (Mla) proteins were also upregulated following RamA overproduction from the pBAD expression vector in *K. pneumoniae* NCTC, but LamB2 is not downregulated (Table S6). These NCTC5055 data were collected during growth in NB, which explains these findings, and confirms the medium dependence of the effects.

Despite these differences in envelope proteome seen during growth of Ecl8Δ*ramR* in MHB and NB, there is little impact of growth media on RamA-mediated envelope permeability reduction. As measured using fluorescent dye accumulation, envelope permeability reduces by ≈75% in Ecl8Δ*ramR* versus Ecl8 during growth in MHB and ≈65% during growth in NB (Fig. 1A).

**Figure 1:**
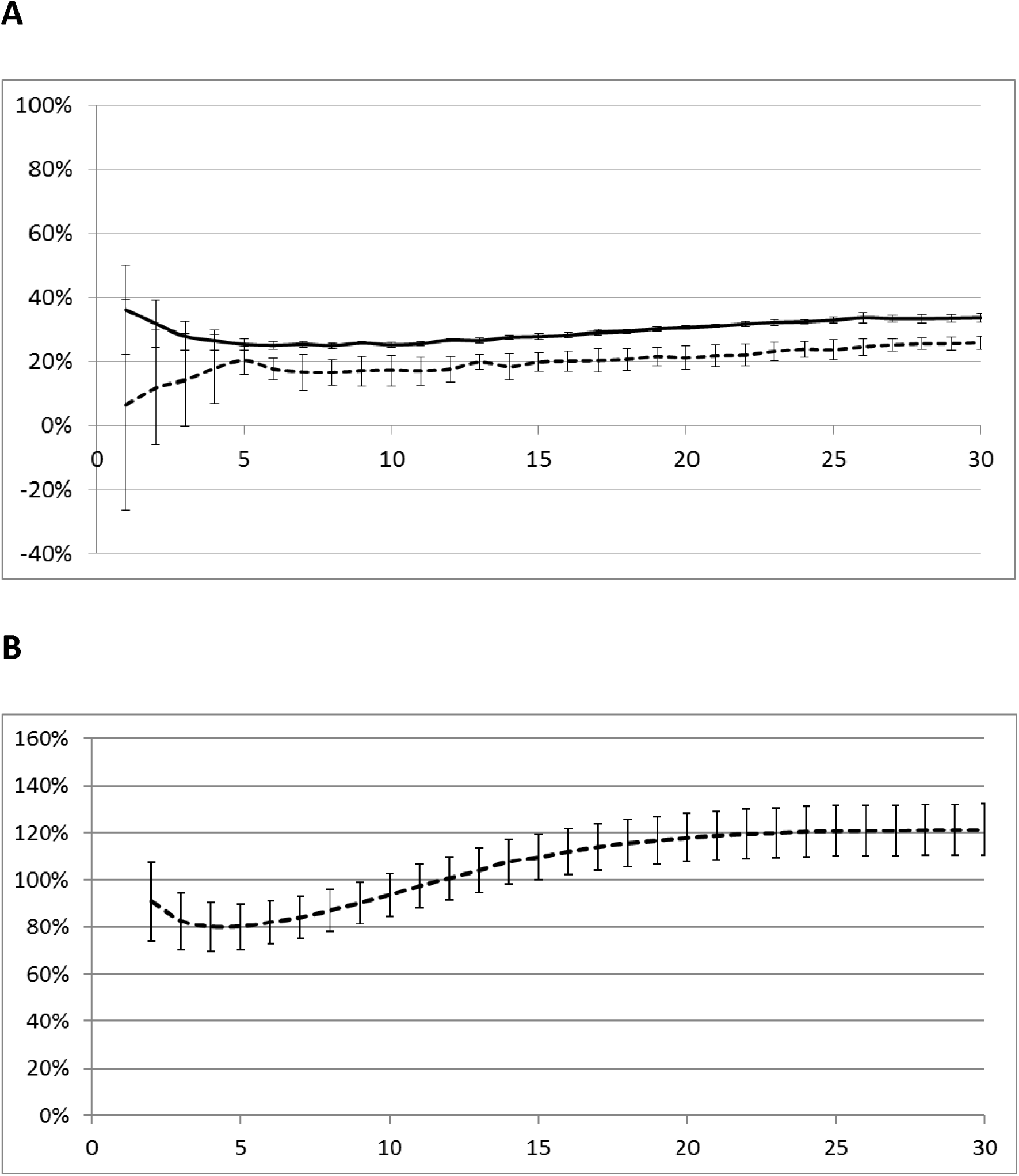
Effect of RamA Over-Production or *micF* Overexpression in *K. pneumoniae* Ecl8 on Envelope Permeability in Different Growth Media. The accumulation of H33342 dye over a 30 cycle (45 minute) incubation period by (A) *K. pneumoniae* Ecl8Δ*ramR* compared with Ecl8 (set to 100%) grown in (solid line) NB and (dashed line) MHB. (B) *K. pneumoniae* Ecl8(*micF*) compared with Ecl8(control) (set to 100%) grown in MHB. Each line shows mean data for three biological replicates with 8 technical replicates in each, and error bars define the standard error of the mean (SEM).

Overall, based on our test of significance, 51 proteins were differentially regulated in NCTC5055 carrying pBAD(*ramA*) versus the pBAD control transformant during growth in the presence of 0.2% w/v arabinose, which stimulates RamA overproduction (Table S7). This is more than the number of proteins differentially regulated in Ecl8Δ*ramR* versus Ecl8 (Tables S5, S6). However, according to Real Time RT-PCR analysis there was 9200 +/− 390 fold (mean +/− SEM, *n*=3) more *ramA* transcript in NCTC5055::pBAD(*ramA*) than in NCTC5055 carrying the pBAD control transformant when grown in the presence of 0.2% (w/v) arabinose, as shown previously.^5^ This is dramatically more than that seen for the “natural” RamA overproducing mutant Ecl8Δ*ramR,* where there is 6.7 +/− 2.2 fold (mean +/− SEM, *n*=3) more *ramA* transcript than in Ecl8, and ≈100-fold more than even the most *ramA* overexpressing clinical isolate in our collection (Table S4). Hence the additional proteomic differences seen in NCTC:pBAD(*ramA*) are likely to be due to spurious occupation of regulatory binding sites by the greatly overproduced RamA, which has previously been reported for *Salmonella* RamA.^2^

### Role of micF in RamA Mediated Control of OmpK35 Levels

There is clear downregulation of OmpK35 porin production following RamA over-production in Ecl8 (Table 2). By analogy with the situation in *E. coli* following over-production of MarA, reduction in OmpK35 levels is likely to be due to transcriptional upregulation of *micF* by RamA. To test whether the short *micF* regulatory RNA can control OmpK35 levels in *K. pneumoniae,* we cloned the *micF* gene, with its own promoter into a high-copy vector pK18^19^ and introduced this recombinant into Ecl8 to boost *micF* transcript levels *in trans.* LC-MS/MS confirmed that OmpK35 levels are downregulated (0.39 fold, *p*=0.005, *n*=3) in Ecl8(*micF*) compared with Ecl8(pK18) during growth in MHB; almost the same downregulation seen in Ecl8Δ*ramR* compared with Ecl8 during growth in MHB (0.43 fold, *p*=0.007, *n*=3 [Table 2]). However, dye accumulation assays revealed *micF* mediated OmpK35 downregulation increases the steady state level of fluorescence so that it is approximately 20% more in Ecl8(*micF*) than in the Ecl8(pK18) control transformant. Surprisingly, therefore, there is a slight increase, not a decrease in overall envelope permeability in the *micF* over-expressing recombinant (Fig 1B). Disc susceptibility testing confirmed that this increase in envelope permeability is sufficient to increase antibiotic inhibition zones against Ecl8(*micF*) (Table S8).

We hypothesised that the reason for this increase in envelope permeability (Fig. 1B) despite OmpK35 levels being reduced is that *K. pneumoniae* responds to reduced OmpK35 levels by downregulating efflux pump production or upregulating porin production to balance envelope permeability. A reciprocal effect: downregulation of OmpK35 in *K. pneumoniae* having reduced AcrAB-TolC-mediated efflux has recently been reported.^30^ In support of our hypothesis, RT-PCR showed that *acrA* levels are reduced in Ecl8(*micF*) versus Ecl8(pK18) (0.53 +/− 0.07 fold [mean +/− SEM, *n*=3]), and this was confirmed by LC/MS-MS to reduce AcrA protein levels (0.62 fold, *p*=0.05, *n*=3). No porin proteins were seen to be upregulated (data not shown). In fact OmpC is downregulated (0.57 fold, *p*=0.02, *n*=3), as is OmpA (0.61 fold, *p*=0.03, *n*=3).

Interestingly, downregulation of *ramA* transcription was seen using Real Time RT-PCR in Ecl8(*micF*) compared with Ecl8(pK18) (0.09 +/− 0.04 fold [mean +/− SEM, *n*=3]). It is possible, therefore, that RamA downregulation may be responsible for the downregulation of AcrA seen in response to *micF-*mediated OmpK35 downregulation. If correct, this is suggestive of a feedback mechanism by which the cell can sense the balance of different factors affecting envelope permeability and control RamA production as necessary.

### Conclusions

RamA overproduction is seen in *K. pneumoniae* clinical isolates,^8^ though a widespread random screen has not previously been used to determine the frequency that this occurs in general. Instead, RamA overproduction has been associated with resistance to specific antimicrobials, e.g. tigecycline.^9^ The data presented here lead us to suggest that RamA overproduction might also be selected in *K. pneumoniae* isolates with acquired β-lactamases, which are very common in the clinic. The effect on carbapenem resistance in *K. pneumoniae* carrying a weakly expressed *bla*_VIM-1_ gene was particularly pronounced, and it was striking to see *ramA* hyperexpression in 3/4 carbapenemase producing bloodstream isolates from a randomly selected collection (Table S4). It was also particularly concerning to see the generation of ertapenem resistance following loss of RamR repressor activity in combination with an AmpC-type enzyme in Ecl8 (Table 1) and in a *ramA* overexpressing clinical isolate carrying CTX-M (Table S4). There may be other ways by which RamA overproduction can be beneficial; for example, it may be a prerequisite for the selection of other resistance-causing mutation events such as target site mutations or those which cause porin loss, e.g. in OmpK36.

LC-MS/MS data presented here show that the primary phenotypic effect of RamA overproduction is mediated by enhanced efflux pump production. There is reduced OmpK35 porin levels, and this can be replicated by overexpressing *micF,* but its effect on antimicrobial susceptibility is low, possibly because of the actions of regulatory feedback cascades. The proteomics methodology we have pioneered for this work has the potential to allow detailed analysis of these and other regulatory cascades in *K. pneumoniae* and other bacteria and may be useful for identifying proteome changes associated with certain resistance phenotypes in clinical isolates and resistant mutants, thereby allowing prediction of resistance profile and mechanism of resistance.

## Acknowledgements

We are grateful to Dr Aisha M. Al-Amri, who made the *bla*_IMP-1_ recombinant plasmid used in this study.

## Transparency Declaration

None to declare – All authors.

